# Reversed graph embedding resolves complex single-cell developmental trajectories

**DOI:** 10.1101/110668

**Authors:** Xiaojie Qiu, Qi Mao, Ying Tang, Li Wang, Raghav Chawla, Hannah Pliner, Cole Trapnell

## Abstract

Organizing single cells along a developmental trajectory has emerged as a powerful tool for understanding how gene regulation governs cell fate decisions. However, learning the structure of complex single-cell trajectories with two or more branches remains a challenging computational problem. We present Monocle 2, which uses reversed graph embedding to reconstruct single-cell trajectories in a fully unsupervised manner. Monocle 2 learns an explicit principal graph to describe the data, greatly improving the robustness and accuracy of its trajectories compared to other algorithms. Monocle 2 uncovered a new, alternative cell fate in what we previously reported to be a linear trajectory for differentiating myoblasts. We also reconstruct branched trajectories for two studies of blood development, and show that loss of function mutations in key lineage transcription factors diverts cells to alternative branches on the a trajectory. Monocle 2 is thus a powerful tool for analyzing cell fate decisions with single-cell genomics.

## Introduction

Most cell state transitions, whether in development, reprogramming, or disease, are characterized by cascades of gene expression changes. Our ability to dissect gene regulatory circuits that orchestrate these changes with bulk cell assays is limited because such experiments report the average of a given population, which may not be representative of any one cell. We recently introduced a bioinformatics technique called “pseudotemporal ordering”, which applies machine learning to single-cell transcriptome sequencing (RNA-Seq) data with the aim to order cells by progression and reconstruct their “trajectory” as they differentiate or undergo some other type of biological transition. Our algorithm, Monocle^1^, places each cell at the proper pseudotemporal position along this trajectory. Subsequently, numerous pseudotime reconstruction algorithms have emerged with varying approaches, capabilities and assumptions.

Despite intense efforts to develop scalable, accurate pseudotime reconstruction algorithms, the current state-of-the-art tools have several major limitations. The first question is how to identify the key genes that mark progress. Most tools either use simple unsupervised schemes that are sensitive to noise and outliers or operate in a “semi-supervised” (and biased) fashion, requiring the user to provide key marker genes, pathways, or information about the experimental design. Second, current algorithms need the user to specify *a priori* the number of cell fates or branch points to expect. Most pseudotime methods can only reconstruct linear trajectories, while others such as Wishbone^2^ or DPT^3^ support branch identification with heuristic procedures, but either require that the user specify the number of branches and cell fates *a priori* or are unable to identify more than one branch in the trajectory. Complicating these matters, few experiments probing complex biological trajectories with multiple fates have been performed to date, and experimentally validating the architecture of these trajectories is extremely challenging, so the accuracy of existing methods remains unclear.

Here, we describe Monocle 2, which applies reversed graph embedding (RGE)^4-6^, a recently developed machine learning strategy, to accurately reconstruct complex single-cell trajectories. Monocle 2 requires no *a priori* information about the genes that characterize the biological process, the number of cell fates or branch points in the trajectory, or the design of the experiment. Monocle 2 outperforms not only its previous version but also more recently developed methods, producing more consistent, robust trajectories. The algorithm scales to experiments with thousands of cells such as those performed with recently developed droplet-based instruments. We apply Monocle 2 to a recent study ^7^ of myelopoiesis to reconstruct the sequence of cell state transitions leading to the granulocyte/monocyte fate decision. We show that Monocle 2 properly diverts cells from mice lacking Gfi1 or Irf8, which are positive regulators of the granulocytic and monocytic fates respectively, onto branches corresponding to opposing fate. Moreover, Monocle 2 correctly positions cells passing through a transient, bipotent state upstream of the branch corresponding to the commitment. These analyses thus experimentally validate Monocle 2’s trajectory reconstruction approach, confirming that it is a powerful, data-driven tool for dissecting cell fate decisions.

## Results

### Unsupervised feature selection identifies key genes that shape developmental trajectories

Single cell trajectories aim to describe the sequence of gene expression dynamics that occur during a biological process such as differentiation. While some biological processes involve the regulation of thousands of genes, others alter only a limited number. A number of studies have identified “ordering genes” on a supervised basis, by comparing groups of cells captured at different times^1^ or by using Gene Ontology annotations^2^, but this imposes strong bias on the shape of the resulting trajectory. Other groups have devised unsupervised selection procedures based on variation across cells^8^, but these techniques in practice can be highly sensitive to user-specified parameters. We therefore sought to design a robust, fully unsupervised procedure for selecting ordering genes for downstream trajectory reconstruction that can scale to thousands of cells.

Our procedure, termed “dpFeature” works by first reducing the dimensionality of the data by projecting the cells onto the top principal components of the data (**Figure 1A**). Next, we further project the data into two dimensions using t-SNE. We then identify clusters of cells using density peak clustering ^9,10^, an efficient, highly scalable one-step clustering algorithm. Finally, we identify the genes differentially expressed between clusters using a GLM-based likelihood ratio test and select the top 1000 significant significant (FDR < 10%) genes as the ordering set.

**Figure 1:**
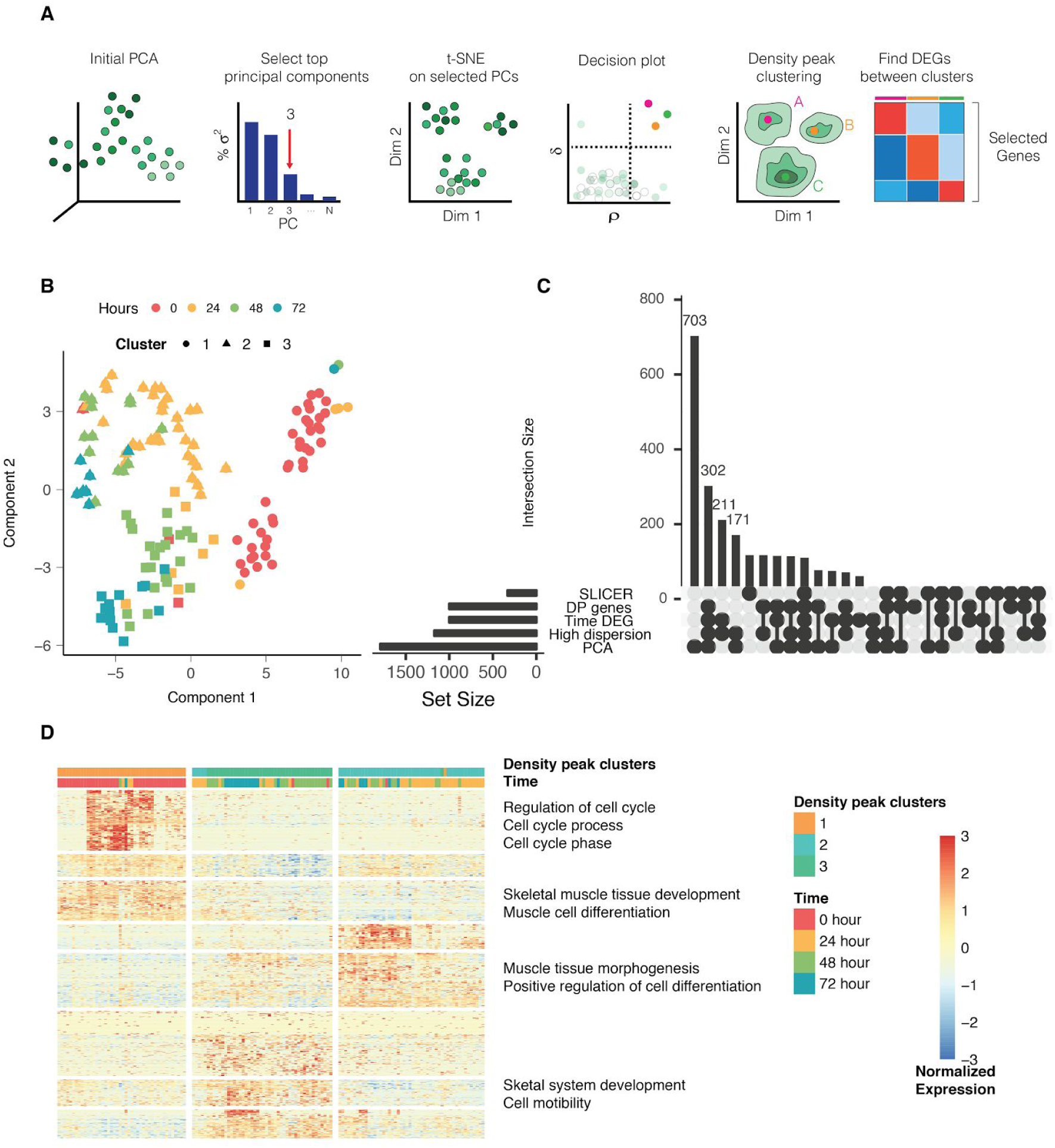
dpFeature detects important genes associated with biological processes. (**A**) Flowchart of unsupervised feature selection based on density peak clustering (dpFeature). The density peak algorithm^9^ is used to cluster cells in a two-dimensional representation generated by t-SNE. Genes that are significantly differentially expressed between clusters are then selected for downstream trajectory inference in Monocle 2. (**B**) tSNE dimension reduction based on top principal components (PCs) and density peak clustering for the HSMM data. (**C**) Five sets of ordering genes are shown: genes that are highly loaded on the first 2 principal components (“PCA”), have high dispersion relative to the mean (“High dispersion”), significantly differ between time points (“Time DEG”), selected via the procedure in SLICER, or identified by dpFeature (“DP genes”). The UpSet plot ^11^ plot shows the number of genes returned by each method (bottom left), along with the size of the possible gene set intersections (up right). (**D**) Clustered heatmap of genes from dpFeature along with selected enriched Gene Ontology terms. Relative transcript counts for each gene (rows) are scaled across cells (columns) and thresholded between the range −3 and 3.

We applied dpFeature to several single-cell RNA-Seq datasets to identify highly relevant genes associated with the biological processes of interest. Firstly, dpFeature was applied to *94* differentiating skeletal myoblast transcriptomes, which were collected over a myoblast differentiation time course. Rather than forming clear, well separated groups of cells with clear margins, the density peak clustering on t-SNE space organized the cells into a more continuous structure with three relatively dense regions of cells (**Figure 1B**). Many of the genes returned by dpFeature were also recovered by supervised comparisons of cells captured at varying times in the experiment and were found amongst the top 1,000 most highly dispersed genes relative to the mean across cells (**Figure 1C**). The dpFeature genes (DP genes) were strongly enriched for Gene Ontology terms associated with myoblast differentiation (**Figure 1C, D**). Applying it to other datasets, including developing lung epithelial cells ^12^, blood monocyte/granulocyte differentiation^7^, and MAR-seq dataset for hematopoiesis ^13^, recovered ordering gene sets with similar levels of overlap with the supervised selection methods (**Figure SI 1**), confirming that dpFeature is a powerful and general unsupervised feature selection approach.

### Reversed graph embedding is an unsupervised approach for learning single-cell trajectories

We next sought to develop a pseudotime trajectory reconstruction algorithm that requires no *a priori* knowledge regarding the number of cell fates or branches. To do so, we employed reversed graph embedding^4–6^, a machine learning technique to learn a parsimonious *principal graph* embedded in the space in which a set of data points reside. A principal graph describes the geometric structure of the data, with every data point distributed around some point on the graph. However, learning a principal graph that describes a population of single-cell RNA-Seq profiles is very challenging because each expressed gene adds an additional dimension to the graph. Learning geometry is dramatically harder in high-dimensional spaces. Reversed graph embedding solves this problem by finding a mapping between the high dimensional gene expression space and a much lower dimensional one and simultaneously learning the structure of the graph in this reduced space. In general, Reversed graph embedding could require a potentially expensive search over a large family of *feasible graphs.* Fortunately, a recent implementation of the reversed graph embedding approach called DDRTree^5^, which limits the set of feasible graphs to trees, performs this search extremely fast even for thousands of cells.

Using DDRTree, Monocle 2 learns a principal graph on a population of single cells, asserting that it describes the sequence of changes to global gene expression levels as a cell progresses through the biological process under study (**Figure 2A**). First, Monocle 2 computes initial coordinates for each cell using using PCA (use of dimensionality reduction techniques such as ICA or diffusion maps is also supported). Next, it learns a tree with an automatically determined number of vertices that “fits” the data set in this space. Then, the cells are moved toward their nearest point on the tree. The algorithm then learns a new tree to fit the cells at their new positions. DDRTree repeats this process until the graph and the cells’ positions have both converged. Once the tree is finalized, Monocle 2 selects a “root” node of the tree as the origin of the trajectory, and it computes the geodesic distance along the graph in the low dimensional space between it and each cell. These distances define the pseudotime of each cell, measuring their progress through the biological process described by the graph. Because Monocle 2 learns a tree structure, in contrast to other methods^1–3,8^, the branch structure automatically emerges. As it updates the cells’ positions and refines the tree, Monocle 2 simplifies the structure of the trajectory, pruning small branches, so that the final trajectory only retains branches that describe significant divergences in cellular states.

**Figure 2:**
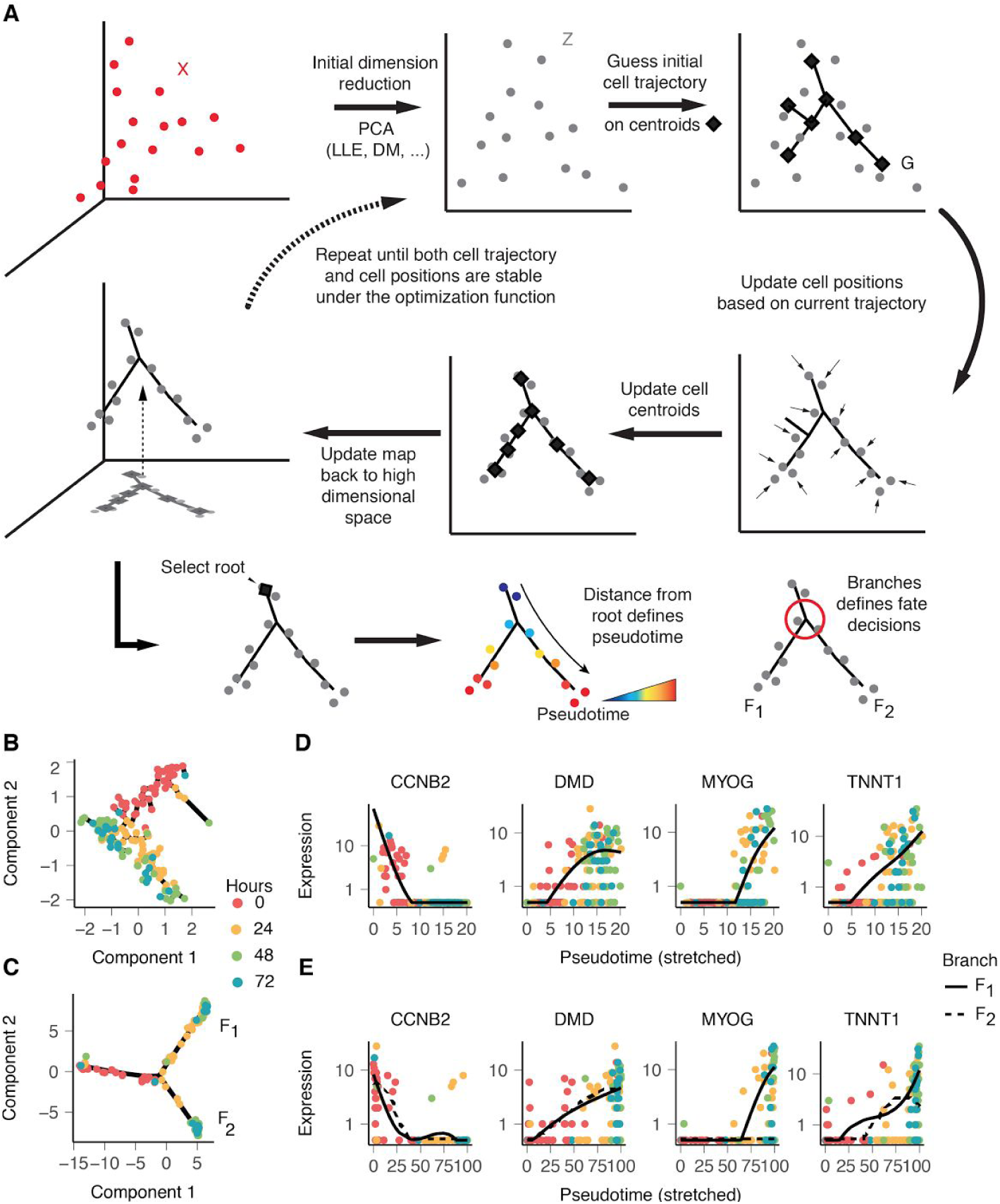
Monocle 2 discovers a cryptic alternative outcome in myoblast differentiation. (**A**) Monocle 2’s approach for learning a single-cell trajectory by reversed graph embedding. Each cell can be represented as a point in a high-dimensional space where each dimension corresponds to the expression level of an ordering gene. The high dimensional data are first projected to a lower dimensional space by any of several dimension reduction methods such as PCA (default), diffusion maps, etc. Monocle 2 then constructs a spanning tree on an automatically selected set of centroids of the data. The algorithm then moves the cells towards their nearest vertex of the tree, updates the positions of the vertices to “fit” the cells, learns a new spanning tree, and iteratively continues this process until the tree and the positions of the cells have converged. Throughout this process, Monocle 2 maintains an invertible map between the high-dimensional space and the low-dimensional one, thus both learning the trajectory and reducing the dimensionality of the data. Once Monocle 2 learns the tree, the user selects a tip as the “root”. Each cell’s pseudotime is calculated as its geodesic distance along the tree to the root, and its branch is automatically assigned based on the principal graph. (**B**) Monocle 1 reconstructs a linear trajectory for the HSMM dataset ^1^. (**C**) Monocle 2 automatically learns the underlying trajectory and detects that cells at 72h are divided into two branches. The same genes selected with dpFeature (Figure 1) are used for ordering for both of Monocle 1 and Monocle 2. (**D**) Kinetic curves for cell cycle marker (CCNB2) and muscle related genes (DMD, MYOG, TNNT1) based on Monocle 1’s ordering. (**E**) Branched kinetic curves for the same markers based on Monocle 2’s ordering.

### Monocle 2 accurately reconstructs single cell trajectories

We first applied Monocle 2 to myoblasts, which we previously reported to differentiate along a linear trajectory^1^. Surprisingly, Monocle 2 reconstructed a trajectory with a single branch leading to two outcomes (**Figure SI 2A, B**, **Figure 2B, C**). Some genes associated with mitogen withdrawal, such as *CCNB2* showed similar kinetics on both branches, but a number of genes required for muscle contraction were strongly activated only on one of the two branches of the Monocle 2 trajectory (**Figure 2D, E**). A global search for genes with significant branch-dependent expression using Branch Expression Analysis Modeling (BEAM) ^14^ revealed that cells along these two outcomes, F_1_ and F_2_, differed in the expression of 664 genes (FDR < 10%), including numerous components of the contractile muscle program. The BEAM analysis suggested that outcome F_1_ represented successful progression to fused myotubes, with high levels of genes required for muscle contraction such as *TNNT2* and *MYH2* as well as key myogenic transcription factors such as *MEF2C* and *MYOG*. In contrast, F_2_ cells failed to upregulate many of these genes (**Figure SI 2**). This is consistent with immunofluorescence measurements of *MYH2,* which show a substantial fraction of isolated nuclei lacking MYH2 and that are not incorporated into myotubes (ref. Figures 1 and 4 of ^1^). Thus, Monocle 2 correctly resolved the two outcomes of myoblast differentiation: successful myotube fusion and failure to fuse.

Although the qualitative dynamics of global gene expression during *in vitro* myogenesis is well established, there is currently no “gold standard” way to order a single-cell dataset. We next simulated single cells undergoing differentiation controlled by a hypothetical gene regulatory network^15,17^. We constructed a set of stochastic differential equations (SDEs) encoding regulatory relationships between 12 regulators important for neuronal development. By combining simulated cells from each of the three possible fates, we simulate a population generated from a branched differentiation trajectory (**Figure 3A**).

**Figure 3:**
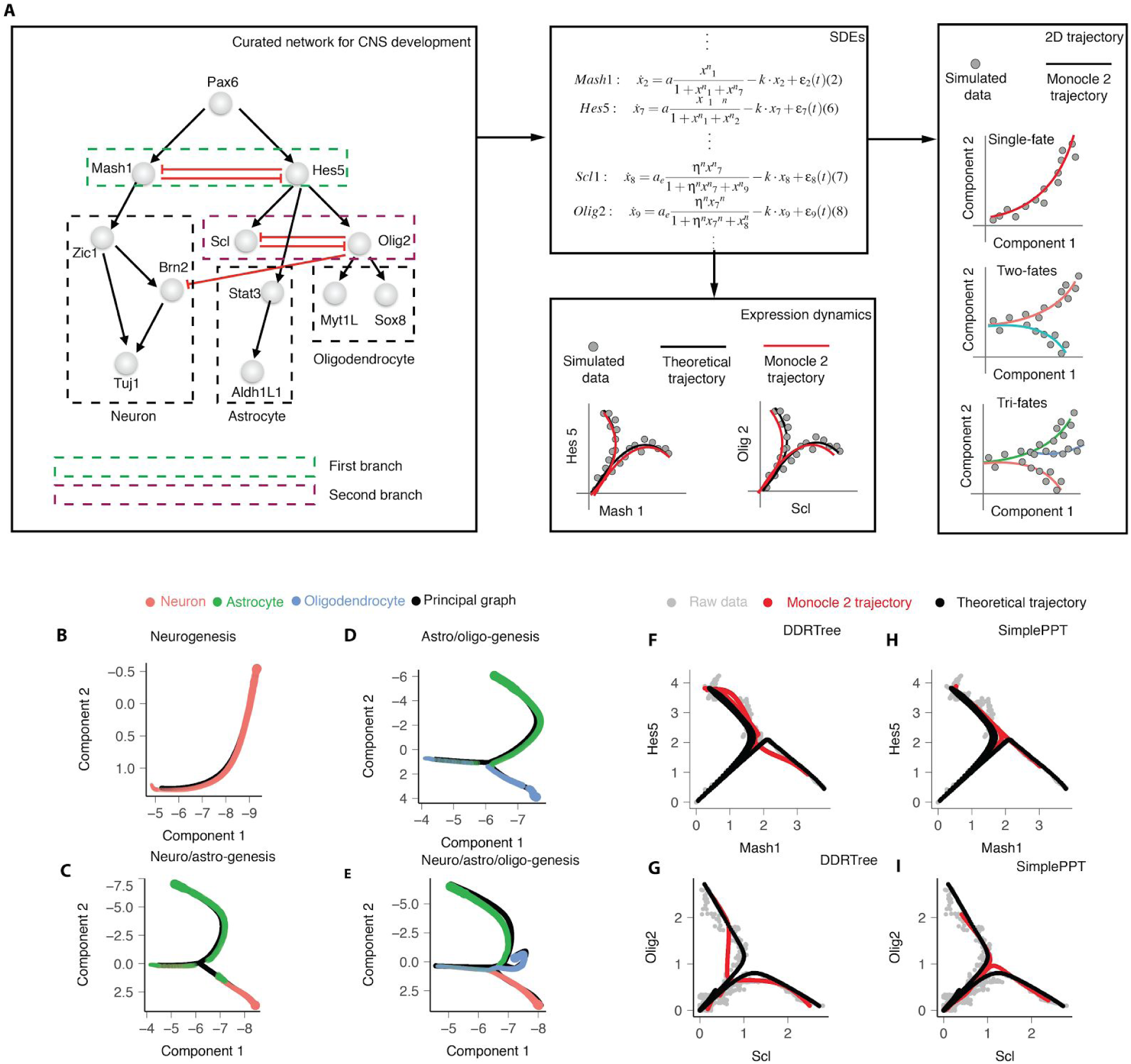
Monocle 2 correctly recovers trajectories driven by simulated gene regulatory networks. (**A**). A hypothetical gene regulatory network and system of stochastic differential equations to drive three-way cell fate specification. The transcriptional regulatory network, modified from Qiu et al, specifies neural progenitor cells to either neurons, astrocytes or oligodendrocytes through a pair of mutual inhibition interactions ^15^. The network summarizes a set of stochastic differential equations that describe gene expression dynamics over time. Initializing this network with small amounts of stochastic noise and following expression kinetics over time simulates the trajectory followed by a single cell, which can be compared to the ideal theoretical trajectory ^16^. (**B-E**) Monocle 2 trajectories learned on four ensembles of simulated data points. (**F-I**) Projecting the principal graph learned by DDRTree into the original gene expression space (**F, G**) or using principal graph from SimplePPT in the same dimension (**H, I**) along with the theoretical trajectory visualizes branching kinetics of individual regulators in the network.

Monocle 2 automatically learned the structure of cells differentiating to one of three distinct fates (**Figure 3B-E**). Our hypothetical regulatory network generates these fates through mutual inhibition between Mash1 and Hes5 and between Scl and Olig2^17^. Monocle 2 reconstructed trajectories that closely corresponded to the most probable path through the high-dimensional gene expression space taken by a cell differentiating to each fate^16^. From the learned trajectories, those genes become mutually exclusive in single cells located at the trajectory’s branch points, suggesting that the trajectory captures mutual inhibition by core regulators of the system (**Figure 3F-I, Figure SI3**). Thus, Monocle 2 correctly learned a trajectory that leads to multiple cell fates, and positioned the branches that give rise to them close to the coordinates at which the genes that govern fate begin to show fate-dependent expression.

### Monocle 2 outperforms alternative reconstruction algorithms

We next sought to compare Monocle 2 to state-of-the art algorithms for inferring single-cell trajectories, including Monocle 1, Wishbone, DPT, and SLICER. Monocle 1 orders cells using a minimum spanning tree built directly on the cells’ coordinates along the top two independent components. Wishbone, which uses a k-nearest-neighbor graph to approximate the cell manifold, assumes a single branch and uses a supervised dimensionality reduction scheme, with “ordering genes” selected based on Gene Ontology^2^. Diffusion pseudotime (DPT)^3^ computes a diffusion map and then infers pseudotime orderings based the diffusion distance between cells. SLICER reduces dimensionality with locally laplacian eigenmaps (LLE) and then constructs a minimum spanning tree on the cells, which yields pseudotimes and branch positions^8^. Unlike Monocle 2, these methods do not construct an explicit, smooth tree. Instead they order cells based on pairwise geodesic distances between them as approximated by a nearest-neighbor graph (Wishbone and SLICER) or minimum spanning tree (Monocle 1) or calculated analytically (DPT). All three methods identify branches implicitly by analyzing patterns in the pseudotime orderings that are inconsistent with a linear trajectory. Furthermore, Wishbone assumes the trajectory has exactly one branch, while DPT can detect more than one, but provides no means of automatically determining how many genuine branches exist in the data. We hypothesized that Monocle 2’s explicit trajectory structure would yield more robust pseudotimes and branch assignments than alternative algorithms.

We tested each algorithm using data from Paul et al, who analyzed transcriptomes of several thousand differentiating blood cells^13^. Monocle 2, DPT, and Wishbone produced qualitatively similar trajectories, with CMP cells residing upstream of a branch at which GMP and erythroid cells diverge (**Figure 4A**). We were unable to run SLICER and Monocle 1 on the full set of 2699 cells, and were forced to downsample the data to 300 cells. From the random downsampled dataset, Monocle 1 constructed a trajectory similar to the reference. SLICER generated a branched trajectory in which the branch occurs within the erythroid population to bifurcate into either CMPs or GMPs. Monocle 2, Wishbone, and DPT produced orderings that were highly correlated with a “reference ordering”, constructed using a panel of markers similar to the approach introduced by Tirosh et al ^18^ (**Figure 4C-E**), while SLICER and Monocle 1 were less so (**Figure 4C-E**). Monocle 2 and Wishbone assigned cells to branches with the highest accuracy, but Monocle 2’s assignments were far more accurate and consistent than Wishbone’s when provided with subsets of the cells (**Figure 4F-H**). Although Monocle 2’s trajectory reconstruction algorithm accepts several user-specified parameters, its results were similar over widely varying values (**Figure SI4**). These benchmarks demonstrate that Monocle 2 produces trajectories that are as accurate and more robust than state-of-art methods and yet makes fewer *a priori* assumptions regarding the number of cell fates generated by the trajectory.

**Figure 4.**
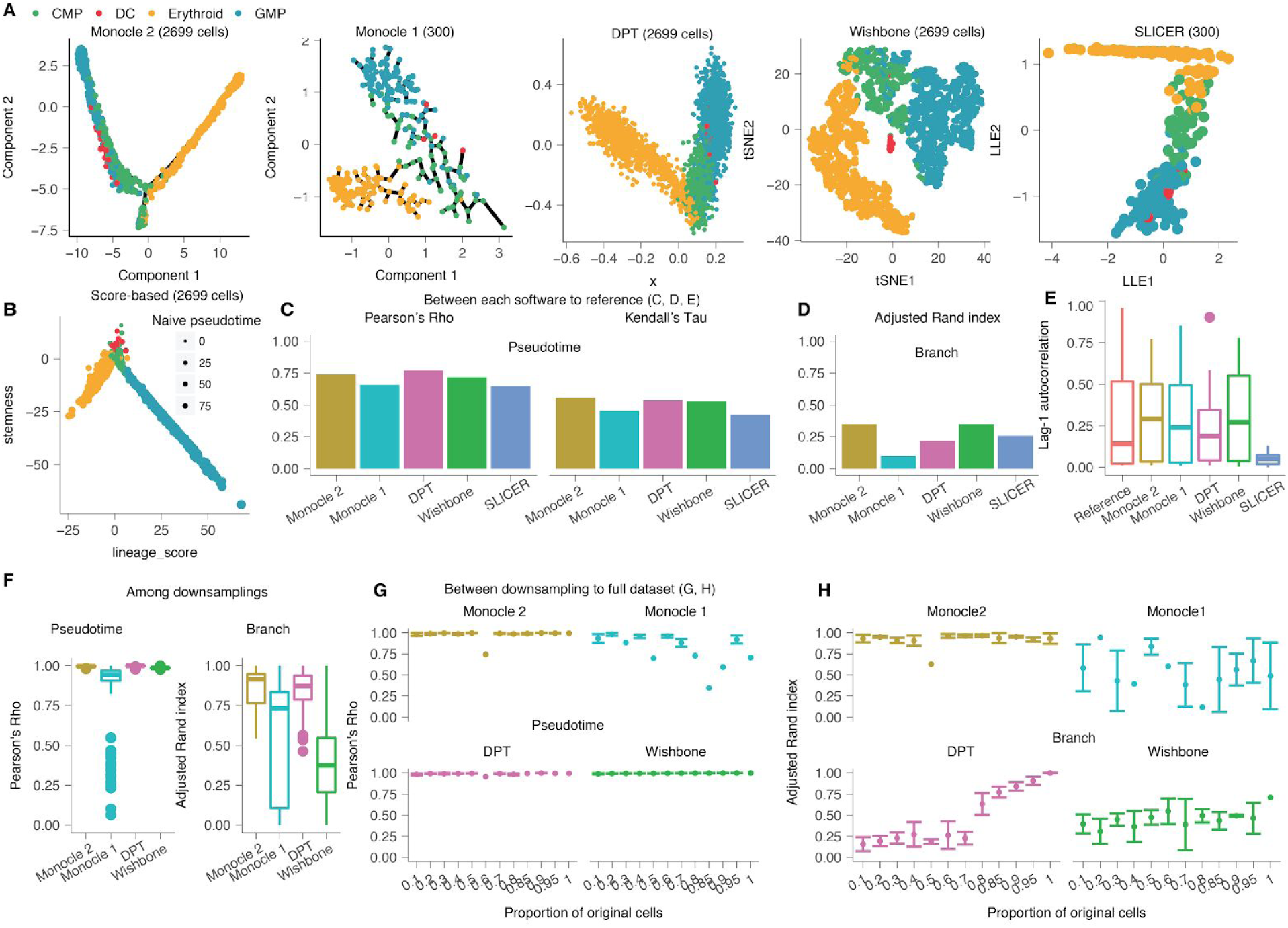
Monocle 2 accurately and robustly reconstructs a bifurcation in hematopoiesis. (**A**) Representation of bifurcation of common myeloid progenitors (CMP) into either erythroid cells or granulocyte-macrophage progenitors (GMP), with trajectories inferred by Monocle 2 and other algorithms. (**B**) Representation of bifurcation of CMP into either erythroid or GMP branch with the score-based “reference trajectory” approach of Tirosh et al ^18^. (**C**) Correlation between the reference trajectory and each tool. (**D**) Adjusted Rand index between the branch assignments from the reference trajectory and branch assignments from each algorithm. (**E**) Lag-1 autocorrelation for the fitted spline curve based on pseudotime from each method for all marker genes used in the score-based ordering. (**F**) Consistency of pseudotime calculation or branch assignments from each algorithm under repeated subsamples of 80% of the cells. (**G-H**) Robustness analysis of pseudotime calculation (**G**) or branch assignment (**H**) for each algorithm run with progressively smaller fractions of the cells. For panels **C-E**, the same 300 random sampled cells are used for all software. In **F-H**, Monocle 2, DPT, and Wishbone all use the full dataset for benchmark while Monocle 1 only uses the same random downsampled 300 cells as above for benchmarking. Only common cells between each pair of downsampling (**F**) or between downsampling and the full dataset (**G-H**) is used for *Pearson’s Rho* or Adjusted Rand index calculations.

We also explored alternative dimensionality reduction and graph learning algorithms. Similar to DDRTree, SimplePPT learns a principal graph on a set of data points, but does not attempt to reduce their dimensionality^4^. The authors of DDRTree and SimplePPT recently reported a new approach, which we refer to as “L1-graph” for learning graphs based on L1-regularized optimization^6^, and could learn cyclical or disjoint trajectories. They also reported a similar strategy, SGL-tree (Principal Graph and Structure Learning for tree)^6^, generalizing SimplePPT, to learn tree structure for large dataset. DDRTree, SimplePPT and SGL-tree reported highly concordant trajectories with PCA, ICA, and diffusion maps (**Figure SI5**). LLE, which is known to be highly sensitive to tuning parameters, sometimes led to incorrect reconstructions with SimplePPT ^19^. L1-graph often reported less refined graphs with numerous minor branches, but captured the overall trajectory structure accurately. The overall approach of reversed graph embedding is thus a powerful, flexible framework for single-cell trajectory reconstruction.

### Monocle 2 captures granulocytic/monocytic specification during blood development

Although Monocle’s trajectories for differentiating myoblasts and common myeloid progenitors were broadly consistent with the known sequence of regulatory events governing those processes, we sought further experimental means of validating the structure of the algorithm’s trajectories. Recently, Olsson *et al* profiled several hundred FACS-sorted cells during various stages of murine myelopoeisis, *i.e.* LSK, CMP, GMP and LKCD34+ cells.We analyzed these cells with Monocle 2 and reconstructed a trajectory with a single major branch and two distinct fates. Lin-/Sca1+/c-Kit+ (LSK) cells were concentrated at one tip of the tree, which we designated the root, with CMP, GMP, and LKCD34+ cells distributed over the remainder of the tree (**Figure 5A**).

**Figure 5.**
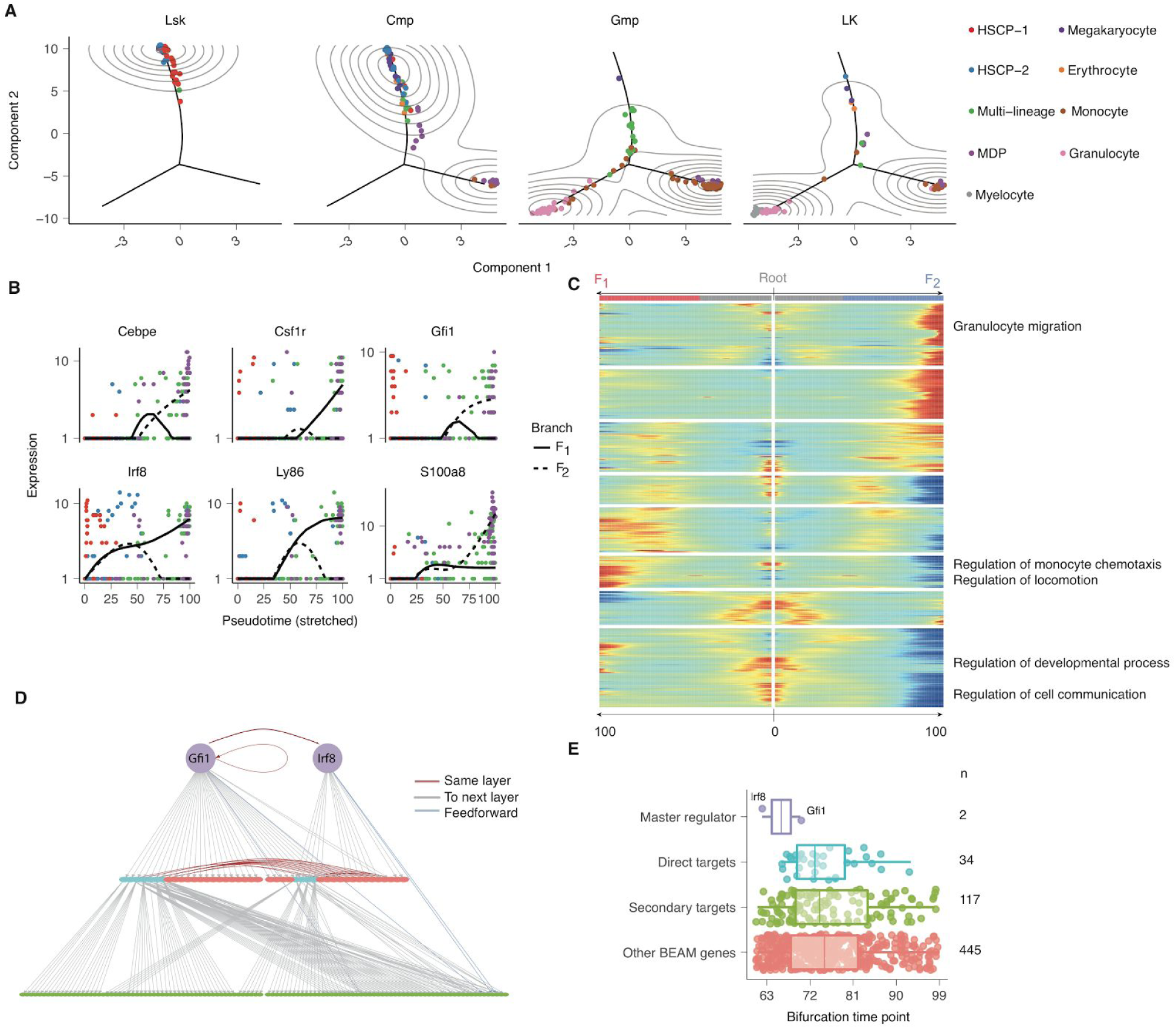
Monocle 2 trajectory branches correspond to developmental fate decisions. (**A**) Monocle 2 trajectory of differentiating blood cells collected by Olsson et al ^7^. Each subpanel corresponds to cells collected from a particular FACS gate in the experiment. Cells are colored according to their classification by the authors of the original study. (**B**) Branch kinetic curves of markers of the monocyte and granulocyte fates. (**C**) Branched heatmap for all the significant branch genes for the wild type data (542 genes, BEAM test, FDR < 10%). (**D**) A network plot describing direct targets of Irf8 and Gfi1 (derived from ChIP-Seq) and secondary targets (derived by motif analysis; see methods). (**E**) Distribution of bifurcation time points for Irf8, Gfi1, and their direct and secondary targets as well as other genes in panel **C**.

Monocle 2 placed cells classified as granulocytes by Olsson *et al* on one of the two branches, while those they deemed to be monocytes were confined to the other. Monocle 2 assigned the CMP cells on the trunk of the trajectory and upstream of the branch. Genes associated with the granulocytic and monocytic programs became progressively more differentially expressed following the branch (**Figure 5B,C**). Many of the genes with significantly branch-dependent expression (BEAM test, FDR < 1%), were bound at their promoters by Irf8 or Gfi1, key activators of the monocytic and granulocytic expression programs, respectively. (**Figure 5D**). These regulators’ expression branched earlier in pseudotime than direct and putative secondary targets (**Figure 5E**), suggesting that the branched trajectory reflects the known regulatory hierarchy of the granulocytic/monocytic fate decision.

In addition to known differentiated cell types, Olsson *et al* detected cells that express a mix of genes specific to different terminal cell fates. They also reported rare, transient cell states that mix hematopoietic/multipotent markers with differentiated markers, which appear mutually exclusive in bulk average measurements of these cell populations. They concluded that both types of “mixed lineage” cells reside in the developmental hierarchy downstream of long-term and short-term HSCs but upstream of cells that have committed to a lineage. Consistent with this interpretation, Monocle 2 positioned mixed-lineage cells and rare transient cells (**Figure 5A, Figure SI6**) just upstream of the the granulocyte-monocyte branch.

### Genetic perturbations divert cells along specific trajectory branches

We next explored how loss of key cell-fate determining genes impacts a developmental trajectory. Adding cells from mice lacking Gfi1 or Irf8 did not substantially alter the structure of the myeloid differentiation trajectory (**Figure 6A**). The tree still contained a single major branch point, with wild-type granulocytes and monocytes confined to distinct branches. However, cells from Gfi1-/- mice were largely excluded from the branch occupied by wild-type granulocytes, and Irf8-/- cells were depleted from the wild-type monocyte branch. That is, the loss of a gene known to activate a fate-specific expression program appeared to divert cells to the opposite fate. Cells from double knockout mice (Gfi1-/- Irf8-/-) were present on both monocyte and granulocyte branches, but concentrated closer to the branch point and away from the tips of the tree, suggesting that they did not fully differentiate.

**Figure 6.**
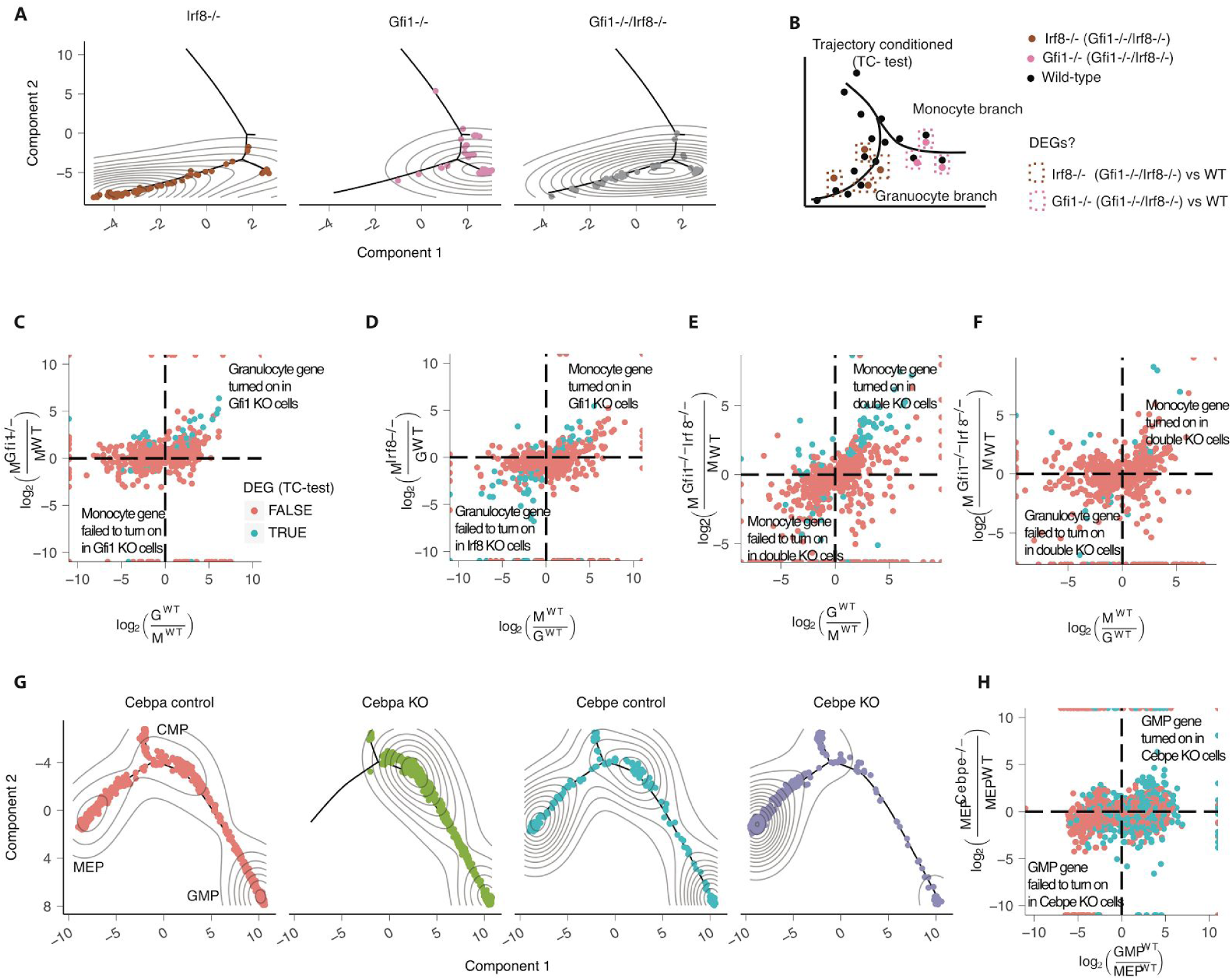
Trajectories reveal how genetic perturbations divert cells to alternative fates. (**A**) Cells with a single knockout of Irf8 or Gfi1 are diverted into the alternative granulocyte or monocyte branch, respectively. Double knockout cells are localized to both granulocyte and monocyte branches but concentrated near the branch point. (**B**) A “trajectory-conditioned” test for identifying genes that are differentially expressed between genotypes that controls for the cells’ positions on the trajectory. (**C-D**) Expression changes for branch-dependent genes (shown in Fig. **5C**) between wild-type granulocytes and monocytes (horizontal axis) plotted against the effects of Gfi1 or Irf8 knockout. The vertical axis in panel **C** shows expression changes in Gfi1 -/monocytes relative to wild-type monocytes, while panel **D** shows changes in Irf8-/- granulocytes compared to wild-type. Genes with significant trajectory-conditioned differences in expression between knockout and wild-type are highlighted (FDR < 10%). (**E-F**) Expression changes in BEAM genes between double KO “monocytes” (**E**) or “granulocytes” (**F**) and wild-type. (**G**) Monocle 2 accurately positions cells from Cebpa-/- or Cebpe-/- collected by Paul *et al.* Loss of Cebpa fully blocked the MEP branch while Cebpe KO partially blocked the GMP branch. (**H**) The trajectory-condition test between Cebpe-/- cells and nearby wild-type cells on the MEP branch reveals aberrant expression of GMP-branch specific genes in MEP branch. Panels (**A-F**) are based on the data from ^<u>7</u>^ while panels (**G, H**) are based on the data from ^13^.

To determine whether Gfi1-/- or Irf8-/-had fully adopted the monocyte and granulocyte expression programs, respectively, we designed a new statistical test similar to BEAM, our recently published method for identifying branch-dependent genes ^14^ This “trajectory-conditioned” test is designed to identify genes that differ between cells from a mutant and the wild type, controlling for where the cells are on the trajectory (**Figure 6B**, **Methods**). Effectively, the trajectory-conditioned test compares cells at similar positions on the trajectory from different genetic backgrounds. Applying it to cells on the monocyte branch revealed that Gfi1-/- cells express higher levels of genes from normally associated with granulocytes than wild-type monocytes (**Figure 6C**). Likewise, cells from Irf8-/- mice on the granulocyte branch showed aberrantly high levels of monocytic genes (**Figure 6B**). Cells from Gfi1-/- Irf8-/- mice recapitulated both these effects, with high levels of monocyte genes in cells on the granulocyte branch and similarly aberrant levels of granulocyte genes on the monocyte branch (**Figure 6E,F**).

Analysis of genetic perturbations from the large-scale transcriptomic study of hematopoiesis reported by Paul *et al* also revealed diversions of cells onto specific branches of the trajectory. Where cells from wild-type mice were distributed along branches corresponding to erythroid (E) and granulocyte/monocyte (GM) fates, cells from *Cebpa-/-* mice were completely excluded from the erythroid branch (**Figure 6G**). Similarly, cells from *Cebpe-/-* were severely depleted from the GM branch, and concentrated at the erythroid branch tip. Similar to diverted cells from Olsson *et al, Cebpe-/-* cells on the erythroid branch showed elevated levels of many GM genes compared to wild-type.

## Discussion

Single-cell RNA-Seq has spurred an explosion of computational methods to infer the precise sequence of gene regulatory events that drive transitions from one cellular state to another. In doing so, pseudotime inference algorithms promise to reveal the gene regulatory architecture that governs cell fate decisions. However, most current methods rely on strong assumptions about the structure of a biological trajectory. Many also require the user to supervise trajectory inference, inject large amounts of *a priori* biological knowledge, or both.

Monocle 2 learns cellular trajectories in a fully data-driven, unsupervised fashion with only limited assumptions regarding its structure. First, Monocle 2 selects the genes that characterize trajectory progress using a simple, unbiased, and highly scalable statistical procedure. It then employs a class of manifold learning algorithms that aim to embed a principal graph amongst the high-dimensional single-cell RNA-seq data. In contrast to previous methods that infer branch structure using heuristic analyses of pairwise distances between cells, Monocle 2 can use this graph to directly identify developmental fate decisions. We have demonstrated through extensive benchmarking that Monocle 2 compares favorably with other tools such as Wishbone without requiring the user to specify the structure of the trajectory.

Analysis of multiple real and synthetic datasets demonstrated that Monocle 2 reconstructs trajectories that faithfully characterize cellular differentiation. Applying Monocle 2 to data from two recently reported studies of hematopoiesis automatically recovered trajectories that captured the erythroid, granulocyte, and monocyte fate decisions. Previously, we showed that loss of interferon signaling can create a new branch in an otherwise linear trajectory that reflects the response of dendritic cells to antigen^14^. Here, we show that cells from mice that lack transcription factors required for establishing specific myeloid fates were diverted onto alternative fates of the same trajectory without altering its structure. Why some loss of function mutations create branches while others divert cells along existing ones is unclear, but this question underscores the increasing power of analyzing single-cell trajectories.

Single-cell trajectory analysis has emerged as a powerful technique for understanding biological processes throughout development. As we dissect more complex communities of cells, the burden placed on trajectory inference algorithms continues to grow. Monocle 2 is a fully unsupervised algorithm that can resolve multiple branch points in a data-driven, unbiased manner. The graph learning approach at the core of Monocle 2 generalizes to trajectories with loops and discontinuities, which will appear in samples containing multiple cell types and proliferative states. Furthermore, because Monocle 2 makes no explicit assumptions about the experiment, it could in principle be applied for pseudospatial reconstruction or other “non-temporal” settings. We also anticipate that Monocle 2 will be useful not just for expression data, but for single-cell chromatin accessibility ^20^, DNA methylation ^21^, or 3D structure ^22,23^ analysis as well. We are confident that Monocle 2 will help reveal how various layers of gene regulation coordinate developmental decision making within individual cells.

## Acknowledgements

We thank I. Tirosh for discussion on marker-based ordering, F. Theis and A. Wolf for discussion on MAR-seq data analysis with DPT, and members of the Trapnell laboratory for comments on the manuscript. This work was supported by US National Institutes of Health (NIH) grant DP2 HD088158. C.T. is partly supported by a Dale. F. Frey Award for Breakthrough Scientists and an Alfred P. Sloan Foundation Research Fellowship.

---

**Supplementary Figure 1. DPFeature shapes the reconstruction of developmental trajectories by selecting informative genes.** (**A**) tSNE plot from dpFeature clusters cells from lung data ^12^ into four different clusters. Color corresponds to sample collection time points while shape corresponds to the cluster assignment. (**B**) UpSetR plot of the ordering genes selected by various procedures, similar to Figure **1C** (**C**) Differentiation trajectories learned with each set of ordering genes. (**D-I**) Similar analysis for the hematopoietic data reported by Olsson et al (panels **D-F**) and Paul et al (panels **G-I**).

**Supplementary Figure 2. Myoblasts differentiate along a branched trajectory.** (**A**) tSNE plot from dpFeature clusters cells from HSMM data into two major clusters. Cells in the upper express a fibroblast-associated gene, *ANPEP,* while the lower cluster has numerous cells expressing muscle-specific genes (MYF5, MYOD1, MYOG). (**B**) Percentage of cells expressing selected muscle or fibroblast-associated genes.(**C**) Branch expression curves for genes in panel **A**. Dashed line indicates the spline fit for cells on the path from the root of the tree in Figure 2D to outcome F_1_, while the solid line indicates the curve for the path to F_2_. (**D**) Distribution of cells detectably expressing *MYOG* along the trajectory. (**E**) Branch kinetic heatmap of significantly branch-dependent genes (BEAM test; FDR < 10%) and selected gene ontology categories enriched in groups of genes with similar kinetics.

**Supplementary Figure 3. Reversed graph embedding accurately learns complex trajectories in simulation datasets.** (**A**) Comparison of the trajectory and branch assignment of learned by several programs (Monocle 2, either with DDRTree or SimplePPT, DPT, SLICER, Monocle 1, and Wishbone) for the simulated two-branch neuro/astro-oligo-genesis process. (**B**) Comparison of each program for learning a complex tree structure.

**Supplementary Figure 4. Effects of parameters of DDRTree on trajectory reconstruction and comparison of Monocle 2 with other state-of-art software** (**A**) A table describes the parameters used in Monocle 2 for trajectory reconstruction. (**B**) Robustness of monocle 2 under a large range of parameters used in DDRTree. Default parameters are used as the reference.

**Supplementary Figure 5. Different Reversed graph embedding techniques reveal the same cryptic myogenesis branch.** (**A**) Trajectory learnt with the different RGE techniques, DDRTree, SimplePPT, SGL-tree, L1-graph on the reduced two dimensional space obtained by running PCA, ICA, LLE or diffusion maps. (**B**) *Pearson’s Rho* or *Kendall’s Tau* correlation between pseudotime calculated with DDRTree initialized with PCA dimension reduction (reference) and pseudotime calculated with DDRTree, SimplePPT, L1 SGL-tree or L1-graph based on reduced dimension space obtained with different dimension reduction methods, using all the dpFeature selected genes as in main figure 1/2. (**C**) *Adjusted Rand index* between clusters calculated with DDRTree initialized with PCA dimension reduction (reference) and clusters calculated with DDRTree, SimplePPT, SGL-tree or L1-graph based on reduced dimension space obtained with different dimension reduction methods, using all the dpFeature selected genes as in Figure **2**.

**Supplementary Figure 6. Monocle 2 correctly positions transient wild-type cells.** (**A**) Gated rare transient cell from ^7^ are enriched near the branch point of monocytes and granulocytes.

## DPfeature: feature selection by detecting DEGs across cluster of cells

An interesting observation we found when analyzing multiple datasets associated with development is that genes informative to differentiation trajectory often form clusters across cell states (Figure 1 of [1], Figure 2-4 of [2] or Figure 2 of [3]). That is, there are clusters of genes which have relative high expression in the progenitor cell states but low expression in the terminal cell state, etc. Inspired by this observation, we developed a simple procedure, termed dpFeature, to automatically select those clusters of genes that have the block expression patterns.

First, dpFeature filters genes that only expressed in a very small percentage of cells (by default, 5%). Second, dpFeature performs PCA on the expressed genes and then users can decide the number of principal components (PCs) used for downstream analysis based on whether or not there is a significant drop in the variance explained at the selected component. These top PCs will be further used to initialize t-SNE which projects the cells into two-dimension t-SNE space. Third, dpFeature uses a recently developed density based clustering algorithm, called density peak clustering [4] to cluster the cells in the two-dimensional t-SNE space. The density peak clustering algorithm calculates each cells local density (*ρ*) and distance (*δ*) of a cell to another cell with higher density. The *ρ* and *δ* values for each cell can be plotted in a so-called decision plot. Cells with high local density that are far away from other cells with high local density correspond to the density peaks. These density peaks nucleate clusters: all other cells will be associated with the nearest density peak cell to form clusters. Finally, we identify genes that differ between the clusters by performing a likelihood ratio test between a negative-binomial generalized linear model that knows the cluster to which each cell is assigned and a model that doesn’t. We then select (by default) the top 1,000 significantly differentially expressed genes, after Benjamini-Hochberg correction, as the ordering genes for the trajectory reconstruction.

## Reversed graph embedding

Monocle 2 uses a technique called reversed graph embedding([5-7] (RGE) to learn a graph structure that describes a single-cell experiment. RGE simultaneously learns a principal graph that represents the cell trajectory, as well as a function that maps points on the trajectory (which is embedded in low dimensions) back to the original high dimensional space. RGE aims to learn both a set of latent points *Ƶ* = (**z**_1_,…, **z**_*N*_} and an undirected graph *𝒢* that connects these latent points. The latent points *Ƶ* in the low-dimensional space corresponds to the input data *𝓧* = {**x**_1_,…, **x**_*N*_} in the high-dimensional space. The graph *𝒢* = (*𝒱*, *𝓔* ) contains a set of vertexes *𝒱* = {*V*_1_,…,*V_N_*} and a set of weighted, undirected edges *𝓔*, where each *V_i_* corresponds to latent point **z**_*i*_, so the graph also resides in the latent, low-dimensional space.

In the context of the single-cell trajectory construction problem, **x**_*i*_ is a vector of the expression values of the *i*^th^ cell in a single-cell RNA-Seq experiment, *𝒢* is the learned trajectory (for example, a tree) along which the cells transit, and **z**_*i*_ is the principal point on *𝒢* corresponding to the cell **x**_*i*_.

RGE learns the graph *𝒢* as well as a function that maps back to the input data space. Let *b_i_*,*_j_* denote the weight of edge (*V_i_*, *V_j_*), which represents the connectivity between z_*i*_ and z_*j*_. In other words, *b_i,j_*> 0 means that edge (*V_i_*, *V_j_*) exists in *𝒢*, and 0 otherwise. Define *f_𝒢_* as the projection function from **z**_*i*_ to some point in the high-dimensional space. To learn *𝒢*, *Ƶ* and *f_𝒢_*, we need to optimize

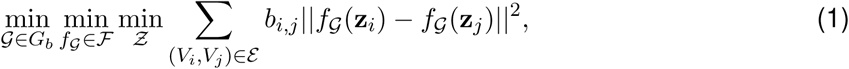

where *G_b_* is a set of feasible graph structures parameterized by {*b_i,j_*, ∀*_i,j_*}, and *𝓕* is a set of functions mapping a latent, low-dimensional point to a point in the original, high-dimensional space.

As shown in [5], the above optimization will learn graph structures in the latent space, but it does not measure the deviations of latent points to the observed data. That is, no effort is made to ensure that the graph nodes are embedded in a way relevant to the cloud of observed data points. To ensure the graph describes the overall structure of the observed data, RGE aims to position the latent points such that their image under the function ***f***_*𝒢*_ (that is, their corresponding positions in the high-dimensional space) will be close to the input data. The optimization problem is formulated as

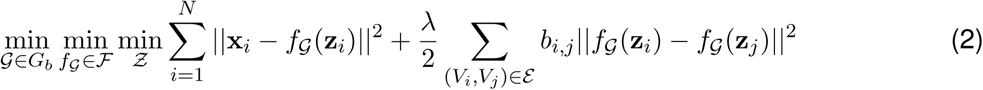

where λ is a parameter that adjusts the relative strength of these two summations. In practice, implementing reversed graph embedding requires that we place some constraints on *𝒢_b_* and *f_𝒢_*, as summarized briefly in the following sections.

## SimplePPT: A simple principal tree algorithm

SimplePPT is the first RGE technique proposed by Mao et al for learning a tree structure to describe a set of observed data points. The tree can be learned in the original space or in some lower dimension retrieved by some dimensionality reduction method such as PCA [5]. SimplePPT makes some choices that simplify the optimization problem. Notably, *f_G_*(*z_i_*) is optimized as one single variable instead of two separate sets of variables. Moreover, the loss function in the reversed graph embedding is replaced by the empirical quantization error, which serves as the measurement between the *f_G_*(*z_i_*) and its corresponding observed points *x_i_*. The joint optimization of *f_G_*(*z_i_*) is efficient from the perspective of optimization with respect to {*b_ij_*}, which is solved by simply finding the minimum spanning tree.

## The principal *𝓛*_1_ graph algorithm

Mao et al later proposed an extension of SimplePPT that can learn arbitrary graphs, rather than just trees, which describe large datasets embedded in the same space as the input [7]. An *𝓛*_1_ graph is a sparse graph which is based on the assumption that each data point (or cell) has a small number of neighborhoods in which the minimum number of points that span a low-dimensional affine subspace [8] passing through that point. In addition, there may exist noise in certain elements of *z_i_* and a natural idea is to estimate the edge weights by tolerating these errors. In general, a sparse solution is more robust and facilitates the consequent identification of test sample (or sequenced single-cell samples). Unlike SimplePPT, this method learns the graph by formulating the optimization as a linear programming problem.

In the same work[7], they also proposed a generalization of SimplePPT, which we term as SGL-tree (Principal Graph and Structure Learning for tree), to learn tree structure for large dataset by similarly considering clustering of data points as in DDRTree. Principal graph and SGL-tree are all treated as SGL in this study.

## DDRTree: Discriminative dimensionality reduction via learning a tree

DDRTree [9], the default RGE technique used by Monocle 2, provides two key features not offered by SimplePPT learning framework. First, DDRTree does not assume the graph resides in the input space, and can reduce its dimensionality while learning the trajectory. Second, it also does not require that there be one node in the graph per data point, which greatly accelerates the algorithm and reduces its memory footprint.

Like SimplePPT, DDRTree learns a latent point for each cell, along with a linear projection function *f_𝒢_* (**z_i_**) = **Wz***_i_*, where **W** = [**w**_1_,…, **w**_*d*_] ∈ *R*^*D×d*^ is a matrix with columns that form an orthogonal basis {**w**_1_,…, **w**_*d*_}. DDRTree simultaneously learns a graph on a second set of latent points 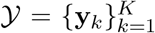. These points are treated as the centroids of 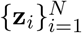 where *K* ≤ *N* and the principal graph is the spanning tree of those centroids. The DDRTree scheme works by optimizing

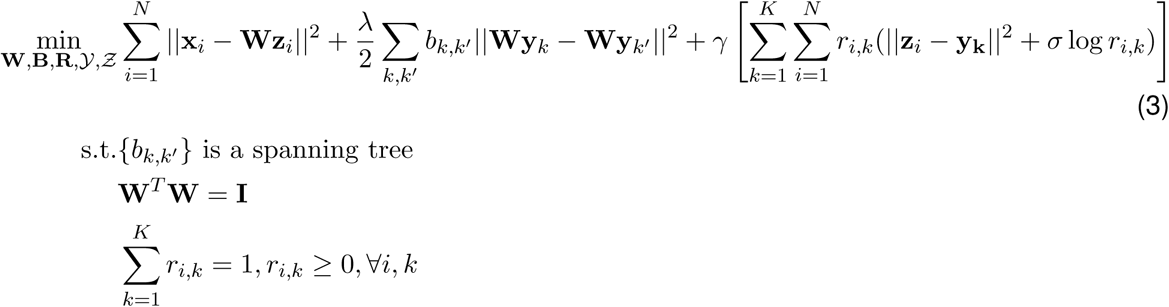

In effect, the algorithm acts as soft *K*-means clustering on points *Ƶ*, and jointly learns a graph on the *K* cluster centers. The matrix R with the (*i*,*k*)th element as *r_i_*,_*k*_ transforms the hard assignments used in *K*-means into soft assignments with *σ* > 0 as a regularization parameter. The above problem contains a number of analytical steps, and can be solved by alternating optimization until convergence. Moreover, because some of the more expensive numerical operations involve matrices that are *K* dimensional (instead of *N* dimensional), they have complexity that is invariant of the size of the input data for a small fixed *K*. In Monocle 2, we provide a procedure to automatically chooses a value of *K* that should work well for a wide range of datasets based on the number of cells *N* in the experiment:

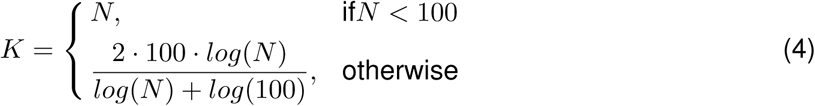

## Initializing RGE with different dimension reduction methods

We implement PCA, ICA, LLE, diffusion maps and pass each function to reduceDimension in Monocle 2 to assess the robustness of different RGE techniques (DDRTree, SimplePPT, SGL) in pseudotime estimation and branch assignment under different dimension reduction initialization approach. PCA is performed with prcomp function from R [10]. ICA is based on *ica_helper* function from Monocle package where *ica_helper* function is built on irlba package [11]. For LLE, we identified the best number of neighbors based on the *select_k* function from SLICER and then run lle function from lle package with the identified *k* where keeping all other parameters as default [12]. For diffusion maps, we run diffusionMap from diffusionMap package [13] with default parameters on the distance matrix of the input normalized expression data. We use HSMM dataset for this benchmark analysis. SimplePPT, DDRTree, SGL are run with default parameters, excepting gamma is set as 0.005 for SGL-tree and 0.1 for *𝓛*_**1**_ graph.

## Pseudotime calculation and branch assignment

Monocle 2 calls DDRTree to learn the principal tree describing a single cell experiment, and then then projects each cell onto its nearest location on the tree. Monocle 2 allows users to conveniently select a tip of the tree as the root and then transverses the tree from the root, computing the geodesic distance of each cell to the root cell, which is taken as its pseudotime, and assign branch segment simultaneously.

DDRTree returns a principal tree of the centroids of cell clusters in low dimension. To calculate pseudotimes, Monocle 2 projects the cells latent points *Z*, to the principal graph formed by principal points, *Y*. For latent points not near principal points internal to the tree, Monocle 2 finds the nearest line segment on the principal tree and then project them to the nearest point on that segment. For latent points near the tip principal points, we will orthogonally project the latent point to the line segment formed by extending the tip principal point and its nearest neighbor principal point in the graph. More formally, we can define a vector between a cell *c* = (*c*_1_,*c*_2_,…), where *c*_1_,*c*_2_,… denotes the coordinates of the cell in the latent space, to the nearest principal point *A* by 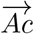. The line segment formed by the two nearest principal points (*A* = (*A*_1_, *A*_2_,…), *B* = (*B*_1_,*B*_2_, …) is 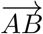. Then we can calculate *t* as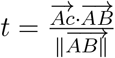. The projection can be calculated as:

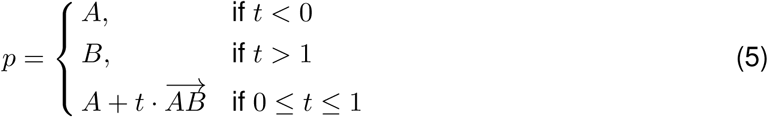

For latent points whose nearest principal points as tips of the principal graph, we will just find the orthogonal projection on the line formed by the tip and the closest point on the principal graph, that is, 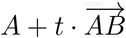.

We then calculate the distance between all the projection points and construct a minimal spanning tree (MST) on the projection points. To avoid zero values of distance between cells projected to the same principal points, which prevents the calculation of a MST, the smallest positive distance between all cell pairs is added to all distance values. This MST is used to assign pseudotime for each cell (See below).

To encode the position of each cell within the branching structure of the trajectory, Monocle 2 performs a depth-first traversal of the principal tree learned during RGE. Without loss of generality, we assume one principal point corresponds to one latent point (for example, in the case we set ncenter = NULL or each cell corresponds to its own cluster). Following the definition introduced in [3], an ordering *π* of cells (principal points) is obtained through a depth first search (DFS) of the learned principal tree starting from the root cell. We can then assign each cell to a trajectory segment, *b_x_*(*G*,*π*,*i*) which specifies the segment, *b*_***x***_ where the cell i is located based on this ordering list, *π*, and the graph structure, *G*. We set *b_x_* = 1 at the root cell and increase a segment counter *b_x_* every time we reach a new branch point. More formally, we can write the formula of segment assignment as:

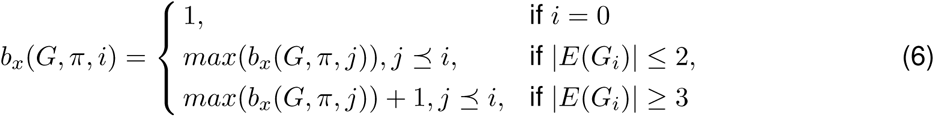

where 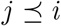 represents the ordering of *i* is precedent of *j*, |*E*(*G_i_*)| represents the degree of cell *i.* For the general cases where the principal points is less than the cell numbers, cells will inherit the segment assignment of their nearest principal point.

Similar to our previous definition of pseudotime [3], Monocle 2 calculates pseudotime based on the geodesic distance of each cell to the root cells on the MST of the projection points. Define pseudotime of cell *i* from a branching biological process *s* with branches given by *b_x_* as *φ_t_*(*b_x_*,*s_i_*), we can calculate its pseudotime recursively by adding the pseudotime of its parent cell on the MST of the projection points (closest cell on the same branch) with the Euclidean distance, 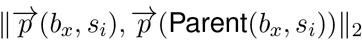, between current and the parent on the MST, by setting the root cell as pseudotime 0. That is,

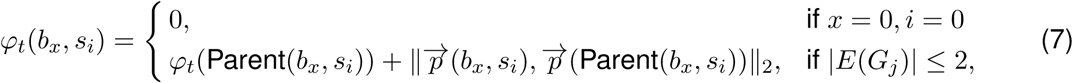

Where 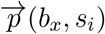 defines the projection coordinates of cell i on the embedded tree.

## Simulating the neuronal differentiation trajectory

Previously, Qiu et al used a system of differential equations describing the dynamics of 12 genes [14] to simulate the hypothetical differentiation process of three cell types from the central nervous system. We use this approach to generate idealized, branched differentiation trajectories for benchmarking Monocle 2. The network mainly consists of two mutual inhibition TF-pairs where the first one (between Mash1 and Hes5) specifies the bifurcation between neuron and glia, the second one (between Scl and Olig2) specifies the bifurcation between astrocyte and oligodendrocyte. All genes are initialized to zero except Pax3. We then follow the gene expression dynamics of all 12 genes by numerically calculating the output of the stochastic differential equation (SDE) for each gene over 400 time steps using the Euler method [15]. Random noise injected into the simulation at each time step leads to the spontaneous, nondeterministic selection of one of three cell fates each time the simulation is run. We therefore run the simulation 200 times to ensure that we obtain a representative path leading from the root to each fate. The gene expression of master regulators (Mash1, Hes5, Scl and Olig2) at the last time step for each simulated developmental trajectory are used to classify trajectories into either neuron, astrocyte or oligodendrocyte type.

We can also obtain a theoretical trajectory from the SDEs as follows. First, we calculate the stable fixed points and metastable ones (saddle points) for the system, which correspond to the progenitor or terminal cell states and the bifurcation points of the differentiation process. According to the A-type integration view of stochastic dynamics, fixed points of their ODE counterparts can be directly regarded as most probable states of SDE [16, 17]. That is, the trajectories (in the sense of dynamical systems) connecting the fixed point corresponding to the progenitor cell state to the other fixed points, corresponding to neuron, glia and oligodendrocyte fate, of the analogous system of ODEs define the most probable path through the SDEs. We take these paths as the theoretical trajectory.

To assess Monocle 2s performance on the simulation data, we selected a simulated trajectory for each of the cell fates, and provided Monocle 2 with the expression values for each of the 12 genes sampled at all time steps. We then run Monocle 2 with default parameters. We also ran Monocle 2 using SimplePPT instead of DDRTree. DDRTree learns the trajectory in the a twodimensional latent space while SimplePPT learns in the original 12 dimension. The trajectories produced by Monocle 2 were then plotted in the high-dimensional space (SimplePPT, Figure 3H-I) or the reversed-embedding dimension one (Figure 3F-G) by projecting low dimension data back to the original dimension (DDRTree) against the theoretical trajectory.

## Comparing different algorithms to a marker-based ordering

In order to test the accuracy of each trajectory reconstruction algorithm, we compared their trajectories to an empirical ordering based on marker genes. Relying on results from Paul et al [1], we first select Pf4, Apoe, Flt3, Cd74 as CMP specific genes, Hba-a2, Car2, Cited4, Klf1 as MEP specific genes and Mpo, Prg2, Prtn3, Ctsg as GMP specific genes. Following the approach of Tirosh et al [18], we then select 100 other genes with expression correlated to these marker genes to calculate a stemness score and GMP or MEP lineage score. We define cells with stemness score larger than 0 as CMP cell and any cells with positive lineage score as MEP cells and negative score as MEP cells. This grouping of cells is later used for branch assignment accuracy evaluation. We then define the reference pseudotime for each cell as:

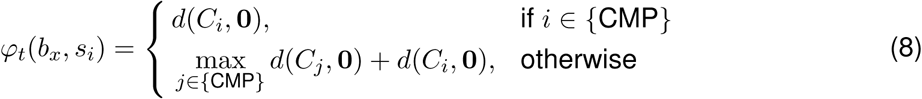

where 0 corresponds to the origin (0,0), *s_i_* corresponds to the stemness score and *l_i_* the lineage score for the lineage to which each cell is assigned, *d*(·,·) represents the Euclidean distance between two points, and {CMP} indicates the set of CMP cells.

We assessed the accuracy of each algorithms trajectory against the reference ordering by two measures of correlation (Pearsons Rho and Kendalls Tau) between their pseudotime values. Pseudotime correlations were computed on the paths from the root to each fate based on the reference ordering separately and then averaged. Since the empirical ordering based on marker genes is not perfect, we also investigate the accuracy of the ordering in terms of the absolute lag-1 autocorrelation of fitted spline curve for the selected marker genes. We first select the trajectory segments corresponding to the transition from the CMP cells to either MEP or GMP cells and then fit a kinetic curve for each marker gene for each transition with a spline curve with three degree of freedom. We then calculate the the absolute lag-1 autocorrelation *r*, which are defined as following:

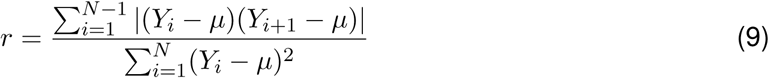

Higher autocorrelation value implies smoother gene expression dynamics based on the ordering.

We used adjusted Rand index (ARI) [19], a common metric used for measuring clustering accuracy, to measure the accuracy of tree segment assignment. Given the number of common cells (300 in this case), denoted as *S*, between the reference ordering and the ordering based on an algorithm (Monocle 2, Monocle 1, DPT, Wishbone or SLICER), and corresponding trajectory segment assignments for reference ordering and ordering based on a different algorithm, *𝒳* and *𝒴*, namely, *𝒳* = {*𝒳*_1_, *𝒳*_2_,…, *𝒳_r_*} and *𝒴* = {*𝒴*_1_, *𝒴*_2_,…, *𝒴*_s_}. The overlap between cells from segment *i* (*𝒳i*) and cells from segment *j* (*𝒴j*) in each of the two orderings is represented by the number *n*_*i*,*j*_ of cells in common, i.e., *n*_*i*,*j*_ ***=*** |*𝒳_i_* ∩ *𝒴*_*j*_|. Define the number of cells with segment *i* from reference ordering is 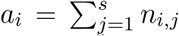, and the number of cells with segment *j* from ordering based on an algorithm is 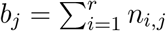. The Adjusted Rand Index is then formulated as

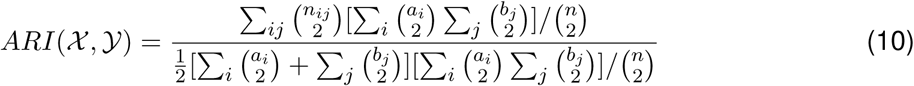

which is a measure of the similarity between two data clusterings (or branch assignment in this case). When ARI is closer to 1, the segment assignment is more consistent between the two orderings.

## Assessing robustness of pseudotime and branch assignments

In order to test robustness of each algorithm in terms of pseudotime estimation and segment assignment, we perform two different downsampling strategies. First, we downsample the MAR-seq dataset [1], selecting 80% of the cells from the full dataset 25 times without replacement. Then we run Monocle 2, Monocle 1, DPT, and Wishbone to construct branched trajectory. SLICER was excluded from the downsampling analysis on account of its long running times and instability on occasional downsample runs. Then we compare all pairs of downsamples by the metrics discussed above.

We also progressively downsample both the MAR-seq dataset [1] over a range of increasing fractions of cells from the full dataset. Sampling is performed without replacement and three different subsets are generated for each proportion to serve as replicates. Then we run each software, including Monocle 1, Monocle 2, DPT, Wishbone, to construct branched trajectories for each fraction, which are compared to the corresponding trajectory built from the full dataset.

ARI is calculated as above while for calculating Pearsons Rho, Kendals Tau we pool all cells for without separating into each different lineage.

## Trajectory-conditioned test

The trajectory-conditioned test is a new type of DEG (differential gene expression) test designed to identify genes that differ between cells from different categories (for example, a mutant and the wild type), controlling for where the cells are on the trajectory. For test in Figure 6, we first identify the range of the pseudotime of the knockout cells on a particular branch. Wild-type cells among this pseudotime range on the same branch are then selected. The knockout cells or the wild-type cells are pooled as two different groups and then a two-group test is performed.

Trajectory-conditioned test can also run by considering both of the pseudotime and the grouping, similarly to the previously defined BEAM test, where the full model is

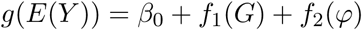

The alternative model is

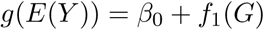

In each of the model, *E*(*Y*) represents the expected value for the transcript counts data *Y* where *Y* is negative binomial distributed or Y *∼ NB*(*r*,*p*)*. g* is a link function; for negative binomial distribution, it is *log. f*_1_(*g*) represents the indicator function for the genotype *G* while *f*_2_(*φ*) represents the non-parameteric function, such as the natural spline implemented in *VGAM* [20] (*sm.ns* function), of pseudotime *φ*..

In practice, we did not use the above formulation because it requires there be enough cells in each category to accurately fit the pseudotime spline curves. However, in situations where both groups are well represented on the trajectory, the above test will likely be more accurate.

## Details on analyzing datasets used in this study

### Analysis of HSMM data

Human skeletal muscle myoblast (HSMM) data from our previous publication [3] is used and converted into transcript counts using Census [21] with default parameters. We first filter cells with total Census counts larger than 1*e*^6^, then filter cells with total Census counts above or below two standard deviation of total Census counts. Potential contaminating fibroblast cells are then identified and removed using a semi-supervised approach, as descibed in the Monocle vignette. We then run dpFeature to select ordering genes (the top 1,000 DEGs are used) on the muscle cells. The top four PC components are used, *ρ* and *δ* are set to 3.8 and 4, respectively based on the decision plot to cluster cells into three clusters using the density peak clustering algorithm [4].

As previously described [21], the ward.D2 clustering method is then applied on the correlation matrix for the row-scaled data (also truncated at 3 or –3) for the Census counts for the 1,000 feature genes between all the genes. To obtain the enriched GO BP terms for the Figure 1D, we perform a hypergeometric test on the corresponding Gene Matrix Transposed file format (GMT) file for each cluster of genes based on the piano package [22]. For PCA based feature selection, we select top 1,000 genes with highest loading from the first two principal components. For SLICER’s feature selection, SLICER is run with default parameters. For high dispersion based feature selection, genes with mean expression larger than 0.1 and empirical dispersion larger than the theoretical dispersion based on empirical mean expression. We used DESeqs method [23] to fit a relationship between mean expression and the variance using a gamma function. For feature selection with DEG test based on known cell group labels, we used the information of collected time point (0 hour, 24 hour, 48 hour or 72 hour) for each cell and identify DEGs between cells from different time points based on generalized linear models and likelihood-ratio test, implemented in Monocle.

To reconstruct the myogenesis trajectory, Monocle 2 is used with default parameters.

### Analysis of lung data

Lung data is processed as described previously [21]. We used transcript counts estimated with spike-in for all analysis of this dataset. We then run dpFeature to select ordering genes (top 1,000 DEGs are used) of the 183 cells. Top five PC components are used, *ρ* and *δ* is set to 3 and 9 based on the decision plot to cluster cells into four clusters. PCA, SLICER, high dispersion based feature selection are done as described above. For DEG based feature selection, time labels of *E*14.5 days, *E* 16.5 days, *E*18.5 days and Adult AT2 cells are used.

### Analysis of MAR-seq data

UMI counts data for the MAR-seq experiment [1] as well as annotation of cells, etc. are downloaded from http://compgenomics.weizmann.ac.il/tanay/?page_id=649. Cells assigned to clusters in the original study are used to classify cells. We combined clusters 1,2,3,4,5,6 as erythroid (1095), clusters 7,8,9,10 as CMP (451), clusters 12,13,14,15,16,17,18 as GMP (1123), cluster 11 as DC (30) and cluster 19 as lymphoid (31), as suggested in the original study [1]. To ensure better comparison with other published work [24, 25], we used all informative genes (3004 genes) identified in the original study [1] for cell ordering. But we also run dpFeature to select ordering genes (top 1,000 DEGs are used) on the transcript counts in Figure SI1. Top five PC components are used, *ρ* and *δ* are set to 3 and 40 based on the decision plot to cluster cells into three clusters. PCA, SLICER, high dispersion based feature selection are done as described above. For DEG based feature selection, cell type labels annotated as above are used. We run Monocle 2 for trajectory reconstruction with default parameters.

Lymphoid cells are removed from our analysis because they belong to a different developmental lineage.

### Analysis of the downsampling dataset with DPT, Wishbone, SLICER

We run DPT (destiny 2.0.3), Wishbone (https://github.com/ManuSetty/wishbone), SLICER (SLICER 0.2.0) using all the same informative genes from the original study [1] as ordering genes. For Monocle 2, we set *norm_method* **=**′ *log*′ which will log transform the expression value (after adding a pseudocount 1, same as below) and then run orderCells function with default parameters. For Monocle 1, we set *norm_method* =′ log′, *reduction_method* **=**′ *ICA*′ in reduceDimension function and set *num_paths* = 2 in the orderCells function. For DPT, we also first log transform the expression value and then run DPT with default parameters. Since Wishbone requires manually selecting the diffusion map components used for trajectory construction which prevent automatic benchmarking, the first two reduced diffusion components from DPT are thus passed into Wishbone (we also tried using the first five diffusion dimension calculated in Wishbone which generally gives worse results). For SLICER, we again first log transform the expression value, then use the *select_k* function to select the best number of neighbors used in lle function. *Min_branch_len* was set to 10 when running *assign_branches* function. Root cells were properly selected for each software to ensure pseudotime calculation for all software starts from the same cells. Both Monocle 2, Monocle 1 and SLICER reduce the high-dimension data into two intrinsic dimensions.

### Analysis of blood data

FPKM values for the blood study is downloaded from synapse (id *syn*4975060). Cell types or gates information are downloaded from the online data along with the original study. We then run Census to estimate the transcript counts. For Figure 5, wild-type cells (excepting the two transition gate cells) are used. We run dpFeature to select ordering genes (top 1,000 DEGs are used) on the Census counts. Top five PC components are used, *ρ* and *δ* are set to 6 and 15 based on the decision plot to cluster cells into three clusters. PCA, SLICER, high dispersion based feature selection are done as described above. For DEG based feature selection, cell type labels based on the original study are used.

We run BEAM to obtain genes significantly branching between granulocyte or monocyte lineages in the wild type blood dataset [26]. Similarly to previously described [21], those BEAM genes are then used to create the branch heatmap in Figure 5C where gene expression pattern along the granulocyte or monocyte branch is clustered into 8 clusters. To obtain the enriched GO BP terms for the Figure 5D, we perform the hypergeometric test on the corresponding Gene Matrix Transposed file format (GMT) file for each cluster of genes based on the piano package [22]. ChIP-seq data for Gfi1 and Irf8 are downloaded and analyzed using MACS2 [27] to identify the potential targets. DHS dataset (*GSE*59992) for the GMP cells are downloaded [28]. We then use FIMO [29] to scan the DHS peaks for the JASPAR vertebrate motif database (version 2016) [30], which give us the regulatory relationship between TFs (with the motif in JASPAR database) and their potential targets. We only focus on those TFs which are both significant BEAM genes and are potentially bound by either Gfi1 or Irf8 based on the ChIP-seq data. We define those TFs as the potential direct targets of master regulator Gfi1 and Irf8. Then we look for the significant BEAM genes which are potentially targeted by those direct targets based on FIMO scanning and define them as the secondary targets. By applying methods discussed previously [21], we calculate the branch time point for all BEAM genes and categorize those genes either as master regulators, direct or secondary targets or other BEAM genes which are not included in all previous sets.

For figure 6, we used the same 1,000 genes selected just from wild-type cells to order the entire dataset, including the wild-type data, the transition gate cells as well as the *Irf*8^-/-^, *GfI*1^-/-^ or *Irf*8^-/-^*Gfi*1^-/-^ cells.

